# Sequence-independent protein domain detection and classification with PRISM

**DOI:** 10.64898/2026.05.22.727336

**Authors:** Abel Tan, Henning Seedorf

## Abstract

The explosion of predicted protein structures has revealed countless novel domain families. However, gold-standard segmentation tools like Chainsaw and Merizo are trained on rapidly obsoleting CATH databases, lack automatic domain classification, and cannot be easily fine-tuned without deep learning expertise. We introduce PRISM, a unified framework enabling sequence-independent, one-shot fine-tuning for simultaneous domain segmentation and classification, bypassing traditional constraints to accurately resolve complex, novel protein architectures.

## Main text

The recent explosion in high-throughput protein structure prediction (Jumper, 2021) has uncovered immense structural diversity, exemplified by the recent discovery of a massively expanded, highly modular family of archaeal adhesin-like proteins (ALPs) (Gupta & Seedorf, 2024). However, efforts to structurally characterize these novel architectures highlight a critical limitation in state-of-the-art domain segmentation tools such as Chainsaw (Wells et al., 2023) and Merizo (Lau et al., 2023). Because these models are anchored to static structural databases like CATH (Sillitoe et al., 2020), they exhibit a strong bias toward well-characterized bacterial and eukaryotic folds, lacking the generalizability to resolve the complex, interleaved architectures of novel families like ALPs. Furthermore, current pipelines decouple domain parsing from functional assignment and offer no practical mechanisms for user-driven fine-tuning on newly discovered domains. To overcome these bottlenecks, we developed PRISM: a unified framework for one-shot domain segmentation and classification. By bypassing sequence-based comparisons, PRISM utilizes a custom-trained object-detection model to visually interpret geometric patterns within Cα-Cα structural distance matrices. This approach successfully resolves architectures that confound standard pipelines, providing an adaptable, high-throughput solution for cutting-edge structural discovery.

PRISM serves as a unified computational framework that integrates sequence-independent domain segmentation and functional classification into a single, high-throughput process. By leveraging a custom-trained YOLOv26 object-detection model, the tool reframes structural parsing as a two-dimensional object detection task, interpreting the distinct geometric configurations found within Cα-Cα distance matrices (Figure 1). This approach eliminates the reliance on sequence alignments and statically biased structural databases, granting the model the flexibility to accurately identify boundaries and automatically label highly complex, uncharacterized protein architectures in one inference step.

**Figure 1:**
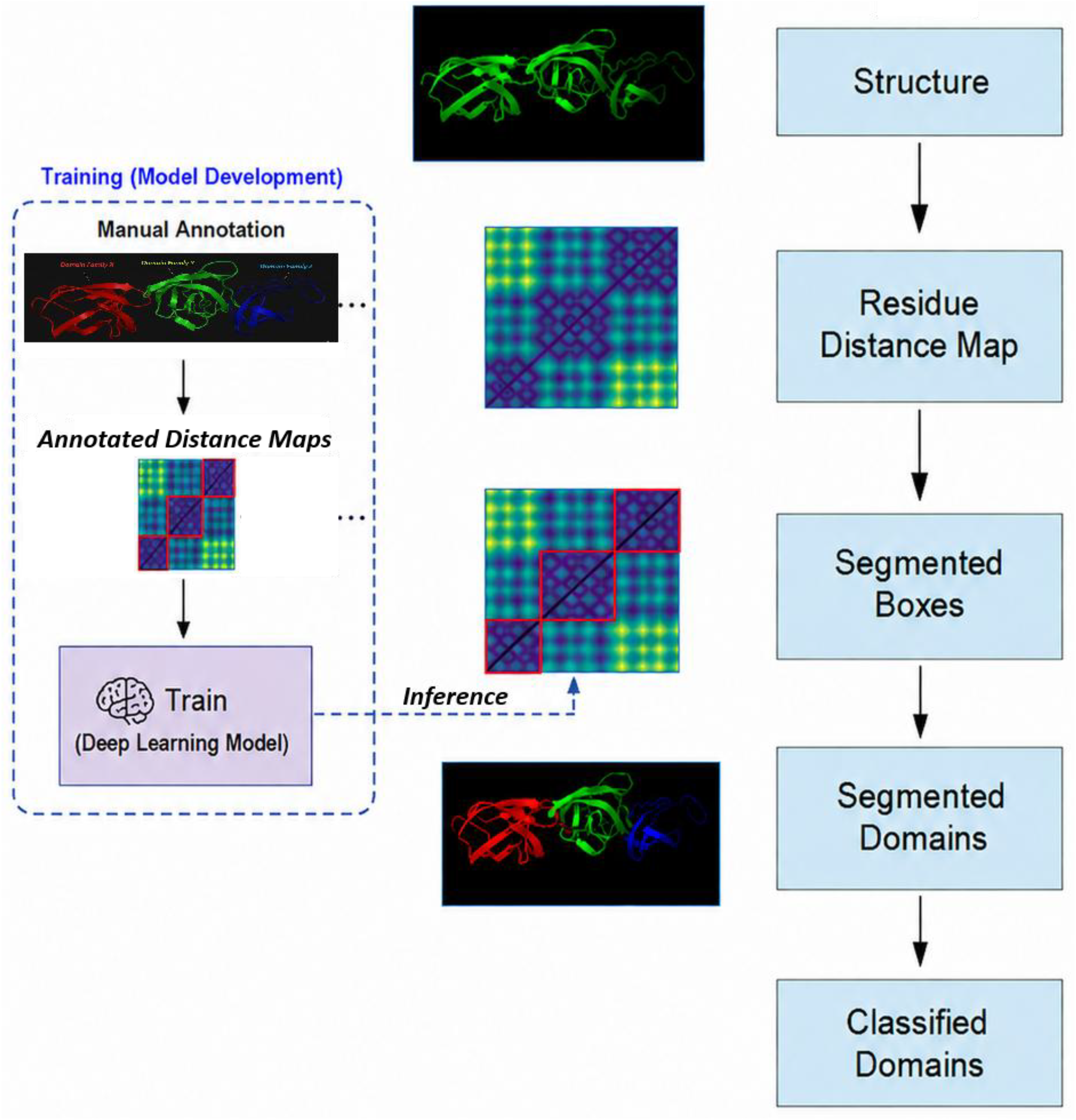
PRISM Pipeline for One-Shot Protein Domain Segmentation and Classification. The framework takes in a protein structure file as input (.pdb or .cif), generates a pairwise Cα-Cα residue distance map, and applies a custom-trained object-detection model to predict bounding boxes along the diagonal. This reframes structural parsing into a 2D visual recognition task, outputting simultaneously segmented and classified domains.

To validate the efficacy of our object-detection approach, we compared the performance of our custom PRISM model against Chainsaw, a state-of-the-art structure-based domain segmentation tool. We trained the PRISM model on a curated dataset of ALP structures, partitioned into strictly non-overlapping training and validation sets to prevent data leakage. For the comparative benchmark, we ran Chainsaw on the corresponding 3D structures. Crucially, to ensure a direct and fair head-to-head comparison, the Chainsaw results were explicitly filtered to evaluate only the specific protein entries comprising the PRISM validation set. To standardize the evaluation metrics across both tools, all domain sub-classifications (e.g., ABD, RBH) were collapsed into a single generic “domain” class.

The custom PRISM model demonstrated superior performance in resolving the novel domain architectures of ALPs compared to existing protein domain segmentation methods. Using a single-class evaluation framework encompassing right-handed β-helical (RBH) domains, archaeal big domains (ABD), and other annotated domains, PRISM achieved the best overall performance on the validation set, with a Precision of 0.948, Recall of 0.892, F1-score of 0.919, and Accuracy of 0.851. In comparison, Chainsaw achieved an F1-score of 0.852 (Precision: 0.852, Recall: 0.806, Accuracy: 0.707), while Merizo showed substantially lower performance, with an F1-score of 0.457 (Precision: 0.549, Recall: 0.384, Accuracy: 0.292) (Table 1). These results support the hypothesis that a custom-trained object detection framework, which visually interprets the characteristic geometric signatures present in protein distance matrices, can be more effective at identifying novel and structurally distinct ALP domain architectures than current state-of-the-art protein domain segmentation tools.

**Table 1:** Performance comparison of domain segmentation methods on the ALP dataset using single-class evaluation across all annotated domain types, including right-handed β-helical (RBH) domains, archaeal big domains (ABD), and other domains. PRISM achieved the highest precision, recall, F1 score, and accuracy compared with Chainsaw and Merizo, demonstrating superior performance for discontinuous domain detection and localization across diverse ALP domain architectures.

| Method | Precision | Recall | F1 | Accuracy |
| --- | --- | --- | --- | --- |
| PRISM | 0.9481 | 0.8923 | 0.9193 | 0.8507 |
| Chainsaw | 0.907 | 0.7879 | 0.8432 | 0.729 |
| Merizo | 0.8706 | 0.7475 | 0.8044 | 0.6727 |

A major limitation of existing protein domain segmentation methods is their inability to simultaneously identify and classify novel domain architectures in a one-shot framework. Current state-of-the-art approaches such as Chainsaw are designed exclusively for segmentation, delineating domain boundaries without assigning functional or structural labels to the detected regions. To address this gap, we evaluated the classification capability of our PRISM framework using a one-shot domain classification task encompassing archaeal big domains (ABD), right-handed β-helical (RBH) domains, and a heterogeneous “other” category representing miscellaneous ALP-associated domains.

The one-shot classifier demonstrated strong performance for the canonical ABD and RBH domain classes, achieving near-perfect discrimination despite the structural diversity of ALPs. ABD domains achieved a precision of 99.4%, recall of 98.1%, and F1 score of 0.99, while RBH domains achieved a precision of 97.6%, perfect recall (100%), and an F1 score of 0.99 (Table 2). Performance for the “other” class was lower (precision: 87.5%, recall: 68.3%, F1: 0.77), which was expected given that this category represents a heterogeneous collection of structurally unrelated domains rather than a single conserved fold. Many “other” regions were conservatively assigned to background, suggesting that the classifier preferentially recognizes strongly conserved geometric signatures while avoiding overclassification of ambiguous regions.

**Table 2:** Class-wise performance metrics for the PRISM one-shot domain classification framework across the three ALP domain categories: archaeal big domains (ABD), right-handed β-helical domains (RBH), and the heterogeneous “other” domain class. Precision, recall, F1 score, and one-vs-rest accuracy were calculated from the confusion matrix. Macro-averaged metrics were computed as the unweighted arithmetic mean across classes to account for class imbalance. The classifier achieved near-perfect performance for the conserved ABD and RBH domain classes, while performance for the “other” category was comparatively lower due to the structural heterogeneity of domains grouped within this class.

| Class | Precision | Recall | F1 Score | Accuracy* |
| --- | --- | --- | --- | --- |
| ABD | 0.994 | 0.981 | 0.988 | 0.971 |
| other | 0.875 | 0.683 | 0.767 | 0.923 |
| RBH | 0.976 | 1 | 0.988 | 0.993 |
| Macro Average | 0.948 | 0.888 | 0.914 | 0.962 |

Despite the heterogeneity of the miscellaneous class, the classifier achieved strong overall macro-averaged performance, with a macro precision of 0.95, recall of 0.89, and F1 score of 0.91. These findings demonstrate that the PRISM framework not only outperforms existing segmentation-only methods in identifying ALP domains, but also uniquely enables simultaneous segmentation and classification of previously unseen domain architectures within a single inference step.

Beyond outperforming automated tools, the model’s sensitivity proved sufficient to correct errors in our previously published manual datasets (Gupta & Seedorf, 2024). We performed a discrepancy analysis between the model’s predictions and the original manual annotations, identifying 16 instances of disagreement. Upon strict visual re-inspection, 12 of these 16 instances (75%) were confirmed to be errors in the original manual annotation, which the model had correctly identified. The majority of these human errors involved “interleaved” domains—specifically, small Repeat-Binding Helix (RBH) domains hidden within long blocks of Archaeal Big Domains (ABDs)—which are visually difficult to distinguish in manual workflows.

Although domain segmentation models such as ChainSaw exist, we found them less suited for the analysed dataset. ChainSaw frequently produced non-contiguous domain predictions, which interfered with our segmentation workflow and lowered model accuracy. In contrast, nearly all ALPs in our dataset contained only contiguous domains, making such predictions biologically inconsistent. Several established domain classification frameworks-including DeepDom (Jiang et al., 2019), DomNet (Yoo et al., 2008), DeepFRI (Gligorijevic et al., 2021), DomSSEA (Rahman et al., 2018), UniDoc (Zhu et al., 2023), DistDom (Mahmud et al., 2022), and Merizo (Lau et al., 2023)-also perform domain segmentation using sequence, structural, or distance-based inputs. However, these models are generally trained or benchmarked on pre-existing structural databases such as CATH, SCOP, ECOD, or CDD, which are inherently biased toward well-characterized bacterial and eukaryotic proteins. As a result, they are limited in identifying novel archaeal architectures such as those present in ALPs. Moreover, most existing frameworks are computationally large and rigid, making fine-tuning impractical without extensive retraining. Given the characteristic features of ALP domain architectures, we therefore opted to train a custom model from scratch. The resulting high segmentation accuracy validates our hypothesis that previously uncharacterized domain architectures can be effectively resolved using a repurposed object-detection model trained directly on structural distance matrices.

The most significant contribution of our approach is the consolidation of domain segmentation and functional classification into a single, high-throughput inference step. Traditional workflows in structural biology typically treat these as decoupled problems: first, tools like Chainsaw or DPAM are employed to delineate structural boundaries, and subsequently, these fragments must be extracted and passed through secondary classifiers or alignment-based searches (such as HMMER or BLAST) to assign functional labels. Our results demonstrate that a repurposed object-detection model effectively bypasses this fragmentation. By training the model to recognize the characteristic “visual signatures” of distance matrices-such as the distinct, dense triangular patterns of ABD domains versus the elongated, repeating helices of RBH domains-the model performs “one-shot” annotation. To our knowledge, no other existing structural bioinformatics tool currently offers this dual capability.

While state-of-the-art tools like Chainsaw are highly effective for well-characterized globular proteins, they encountered significant challenges with the unique modularity of ALPs. The high failure rate (4.4%) observed with Chainsaw, primarily due to GPU memory constraints, highlights a bottleneck for structure-based algorithms when dealing with exceptionally large proteins (often several thousand amino acids in length). Our PRISM model, by operating on a fixed-resolution image representation of the distance matrix, remains computationally lean and robust, successfully annotating the entire 1,077-protein dataset without technical failure.

The discrepancy analysis between our model and previous manual annotations (Gupta & Seedorf, 2024) reveals a critical advantage of machine learning over human expert inspection. This precision is vital for the accurate evolutionary classification of archaeal proteins, where a single missed domain can misrepresent a protein’s entire group architecture. Together, these findings highlight the potential of image-based object-detection frameworks to overcome key limitations of conventional domain annotation pipelines and enable scalable discovery of previously unresolved protein architectures.

## Notes

### Competing Interest Statement

The authors have declared no competing interest.

